# Integrative taxonomy reveals cryptic diversity in Chilean Trichomycterinae (Siluriformes, Trichomycteridae)

**DOI:** 10.64898/2026.07.26.736611

**Authors:** Claudio Quezada-Romegialli, Gloria Arratia

## Abstract

Species-level delimitation within the genus *Trichomycterus* remains one of the main systematic challenges within the Trichomycterinae, particularly in lineages characterised by conservative external morphology, high apparent intraspecific variation, and historical diagnostic criteria based primarily on body proportions, colouration and a few meristic characters. In central Chile, *Trichomycterus areolatus* has traditionally been interpreted as a widely distributed and morphologically variable species, whilst *T. maculatus*, originally described from “Santiago du Chili”, has remained subordinate to this broad conception without a modern phylogenetic reassessment. Here we reassess the specific boundaries of *T. areolatus sensu lato* using an integrative approach that combines complete mitogenomes, estimates of genetic divergence and comparative morphology of the cephalic laterosensory system associated with the neurocranium. Phylogenetic analyses reveal *T. areolatus sensu lato* to be non-monophyletic and identify a deeply divergent lineage, geographically coherent and attributable to *T. maculatus*. This lineage differs from restricted *T. areolatus* by extensive mitochondrial divergence, comparable to that observed between recognised species of Trichomycterinae, and by discrete characters of the cephalic lateral line system, primarily related to the continuity of the supraorbital canal and the arrangement of the associated pores. The congruence between mitogenomic, nuclear and morphological evidence supports the revalidation of *Trichomycterus maculatus* Valenciennes, 1846, and calls for a more restricted geographic circumscription of *T. areolatus*. These results demonstrate that the diversity of Trichomycterinae in central Chile has been underestimated, modify previous interpretation of the distribution of the species involved, and highlight the value of integrating mitogenomics and neurocranial/laterosensory characters into the taxonomy of morphologically conservative siluriform lineages.

## 1. Introduction

Neotropical freshwaters harbour the world’s richest freshwater fish fauna, yet the species limits and phylogenetic relationships of many lineages remain incompletely resolved (Reis, Lecointre, & De Pinna, 2025). Among these lineages, Trichomycteridae (Teleostei: Siluriformes) comprises approximately 473 valid species distributed from Costa Rica to Patagonia and on both sides of the Andes (Fricke, Eschmeyer, & Fong, 2026). The family occupies a broad range of aquatic environments, from lowland rivers to high-altitude Andean streams and lakes above 4,500 m a.s.l. (Arratia F., 1983; Arratia F., Chang G., Marque, & Rojas M., 1978; G. Arratia & Menu-Marque, 1981). Despite their ecological and morphological importance, trichomycterids pose substantial systematic challenges because many new species have been described on the basis of labile morphological characters, while numerous taxa exhibit high levels of intraspecific morphological variation (Ochoa et al., 2020, 2017). These complexities have resulted in repeated changes to their taxonomic classification and persistent disagreement regarding the monophyly and composition of major subfamilies (Reis et al., 2025).

Within Trichomycteridae, the subfamily Trichomycterinae accounts for most of the species diversity of the family and represents one of the major systematic challenges in the taxonomy of Neotropical and Austral South American catfishes (Fernandez, Arroyave, & Schaefer, 2021; Ochoa et al., 2017). Classical morphological revisions had already revealed difficulties in delimiting the subfamily and its genera based on osteological, laterosensory, and external characters, particularly because some lineages exhibit combinations of generalized character states or characters of uncertain polarity (G. Arratia, 1990, 1998; Henschel, Mattos, Katz, & Costa, 2018). In this context, *Trichomycterus*, the most species-rich genus within Trichomycterinae, has historically been treated as a problematic taxonomic assemblage, diagnosed more by the absence of derived characters defining other genera than by exclusive synapomorphies (G. Arratia, 1990). This condition, together with a relatively conserved external morphology of limited diagnostic value for species delimitation, is consistent with the recovery of *Trichomycterus sensu lato* as non-monophyletic in recent molecular analyses (Fernandez et al., 2021; Henschel et al., 2018; Ochoa et al., 2017).

A remarkable example of this taxonomic complexity is *Trichomycterus areolatus* (Valenciennes, 1846), one of the most widely distributed obligate freshwater fishes in Chile, with a range extending approximately 1,400 km from River Huasco to Chiloé Island (Barber, Unmack, Pérez-Losada, Johnson, & Crandall, 2011; Unmack, Bennin, Habit, Victoriano, & Johnson, 2009). Traditionally interpreted as a single species exhibiting substantial morphological variation across its geographic range, *T. areolatus* has remained a problematic taxon for which alternative species hypotheses have never been formally evaluated using integrated molecular and morphological approaches. However, recent phylogenetic studies based on molecular data have begun to reveal cryptic diversity within morphologically conserved trichomycterine taxa (Barber et al., 2011; Unmack et al., 2009). In a phylogeographic study spanning the species’ entire distribution, Unmack et al. (2009) analysed the mitochondrial cytochrome b gene in Chilean populations of *T. areolatus*, using *Bullockia maldonadoi* (Eigenmann, 1919) to root the tree, and recovered seven genetically distinct and well-supported clades (Figure 1a), with no substantial geographic overlap. These seven clades exhibited model-corrected mitochondrial divergences ranging from 1.0% to 9.1%, values comparable to those observed between recognised sisters species in other fish groups, suggesting that each clade could represent an evolutionarily distinct entity under phylogeographic or character-based species concepts. To evaluate these molecular findings, Barber et al. (2011) expanded the original analysis by adding two nuclear markers (rag1 and gh) to the mitochondrial dataset for *T. areolatus* previously analysed by Unmack et al. (2009). The results of this second study again confirmed that *T. areolatus sensu lato* was structured into several genetic groups (Figure 1b), showing patterns of intraspecific differentiation similar to those recovered previously, although with lower resolution at basal nodes because of the more conserved nature of some nuclear markers and without using *B. maldonadoi* as outgroup. These genetic groups were interpreted as “evolutionarily significant units” under multiple species-delimitation criteria. However, the explicit taxonomic implications of these findings were neither rigorously examined nor integrated with morphological analyses.

**FIGURE 1.**
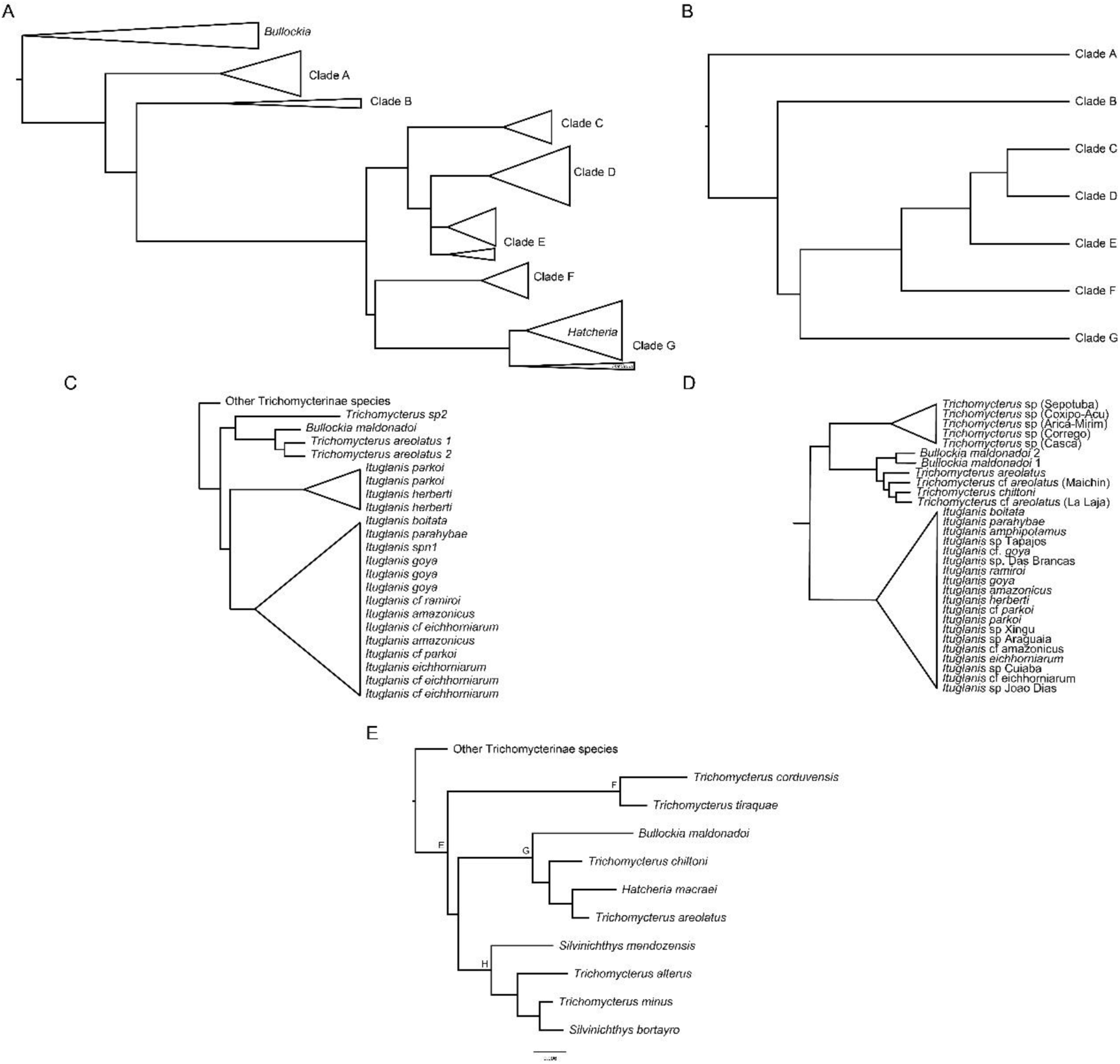
Alternative phylogenetic hypotheses involving *Trichomycterus areolatus* and closely related trichomycterines. (a) Mitochondrial phylogeographic hypothesis showing the main geographic clades recovered within *T. areolatus* sensu lato according to Unmack et al. (2009). (b) Simplified mito-nuclear hypothesis for the principal *T. areolatus* clades according to Barber et al (2011). (c) Multilocus hypothesis including *Bullockia maldonadoi*, *T. areolatus* and *Ituglanis* according to Ochoa et al (2017). (d) Phylogenomic UCE hypothesis showing the position of *T. areolatus* within the areolatus clade relative to *B. maldonadoi*, *T. chiltoni* and *Ituglanis* (Ochoa et al., 2020). (e) Multigene hypothesis of Fernandez et al. (2021) for Trichomycterinae, recovering *T. areolatus* in association with southern Andean trichomycterines, including *Bullockia*, *Hatcheria* and *T. chiltoni*. Topologies were redrawn and simplified from the original studies to emphasize the phylogenetic context of *T. areolatus*; branch lengths are not comparable among panels.

Subsequent phylogenetic analyses conducted by Ochoa et al. (2017; Figure 1c), using multilocus molecular data, placed *T. areolatus* in close relationship with *B. maldonadoi*, indicating that the evolutionary affinities of this taxon were not consistent with a straightforward placement within *Trichomycterus sensu lato*. Later, the phylogenomic analyses of Ochoa et al. (2020, their Figure 4; Figure 1d), based on ultraconserved elements, explicitly recovered an “areolatus clade” comprising *B. maldonadoi*, *T. areolatus*, *T.* cf. *areolatus* from Maichín, *T. chiltoni*, and *T.* cf. *areolatus* from La Laja. This configuration suggested that *T. areolatus* sensu lato may represent a paraphyletic or non-monophyletic assemblage, in which lineages tentatively assigned to *T.* cf. *areolatus* are associated with other nominal southern Andean taxa. This pattern was subsequently expanded by Fernández et al. (2021), who, after incorporating *Hatcheria macraei* (Girard, 1855), species of *Silvinichthys*, and a broader representation of Andean *Trichomycterus* species, recovered *Silvinichthys* and two Argentinean *Trichomycterus* species as the sister group to the clade comprising *B. maldonadoi*, *T. chiltoni*, *H. macraei*, and *T. areolatus* (Figure 1e).

Taken together, the available evidence indicates that *T. areolatus sensu lato* does not simply represent a widely distributed species exhibiting pronounced geographic variation, but rather a complex of lineages associated with the southern Andean clade of Trichomycterinae. However, most of this evidence derives from phylogeographic or phylogenetic studies designed to resolve broad-scale diversification patterns rather than to formally assess the taxonomic limits of Chilean populations assigned to *T. areolatus*. Consequently, a gap remains between the molecular identification of divergent lineages, the phylogenetic interpretation of their non-monophyly, and the morphological evaluation required to support explicit taxonomic decisions.

This gap is particularly relevant in central Chile, where the historical localities associated with two species described by Valenciennes (1846) are located: *T. areolatus*, from the “río de Santiago”, and *T. maculatus*, from “Santiago du Chili”. Both descriptions were brief and based primarily on general morphological comparisons, without detailed diagnoses that would allow their species boundaries to be assessed precisely. The taxonomic situation subsequently became even more complex when Philippi (1866) described three additional species from the same river, *T. marmoratus*, *T. pallens*, and *T. tigrinus*, whose type specimens have never been located. This instability was addressed by Arratia and Chang (1975), who, after conducting a detailed osteological examination of more than 400 specimens, concluded that the skeletons of the two species were indistinguishable and synonymised *T. maculatus* with *T. areolatus*. This conclusion must be interpreted in the context of the methodological tools available at the time. The analysis of Arratia and Chang (1975) focused on general osteological characters of the neurocranium and postcranial skeleton, at a time when the morphology of the lateral-line system and skin had not yet been formalised as a detailed comparative framework for diplomystids and early-diverging loricarioids, as was later accomplished by Arratia and Huaquín (1995). Therefore, the proposed synonymy represented a valid conclusion under the osteological standards of the time, but did not necessarily exclude discrete differences in specialised components of the cranial laterosensory system, whose systematic relevance was not extensively explored until 1995 by the same author.

In this context, integrating complete mitochondrial genomes with comparative morphological characters, particularly those of the cranial laterosensory canal system, provides an opportunity to reassess species boundaries within the *T. areolatus* complex. This approach is especially appropriate in Trichomycterinae, where external morphology may be conserved or ambiguous, whereas certain osteological and laterosensory characters may provide independent diagnostic evidence for recognising evolutionarily differentiated lineages.

In this study, we test the monophyly of *T. areolatus sensu lato* in central Chile and reassess the taxonomic status of *T. maculatus* using an integrative approach based on complete mitochondrial genomes and comparative morphology of the cranial laterosensory canal system. Specifically, our objectives are to: (1) infer mitogenomic phylogenetic relationships among Chilean trichomycterines, with emphasis on the lineages currently assigned to *T. areolatus*; (2) quantify genetic divergence among the principal recovered lineages; (3) compare the configuration of the cranial laterosensory system between the focal taxa; and (4) evaluate whether congruence between molecular and morphological evidence supports the non-monophyly of *T. areolatus sensu lato* and the revalidation of *T. maculatus* as a distinct species.

## 2. Materials and Methods

### 2.1 Next-gen sequencing, assembly, and annotation of newly generated mitogenomes

To infer mitogenomic phylogenetic relationships among Chilean trichomycterines, specimens provisionally assigned to *T. areolatus* and *T. maculatus* were obtained from the Maipo River basin, the type locality of both species described by Valenciennes (1846). In addition, *T. chiltoni* and *B. maldonadoi* were included from Nonguén Stream, the type locality of both taxa, together with *Hatcheria macraei* from the Tunuyán River, Argentina, an undescribed species of *Trichomycterus* from the Chilean Altiplano, and *Nematogenys inermis* (Guichenot, 1848) from its type locality in the River Maipo. Genomic DNA was extracted using the DNeasy Blood & Tissue Kit (Qiagen), following the manufacturer’s instructions. Sequencing libraries were prepared using the MGIEasy Fast FS Library Prep Set V2.0 (MGI, China), including enzymatic fragmentation, adapter ligation, indexing, and amplification. High-throughput sequencing was performed on a DNBSEQ-G400 platform (MGI, China) using the DNBSEQ-G400RS High-throughput Sequencing Set and obtaining 150-bp paired-end reads. Library preparation and sequencing were conducted by TCL Genomics, Santiago, Chile.

Raw paired-end reads were processed using a standardised workflow implemented in custom Bash scripts. Forward and reverse FASTQ files were first evaluated with FastQC (Andrews, 2010) to characterise per-base sequence quality, GC content, sequence-length distributions, and potential adapter contamination. Adapter removal and quality filtering were subsequently performed with Cutadapt (Martin, 2011) in paired-end mode, removing platform-specific MGI adapters. Reads were quality-trimmed at both ends using a Phred threshold of Q20, and reads shorter than 50 bp after trimming were discarded. Only read pairs in which both reads met the specified thresholds were retained.

Following quality filtering, mitogenomes were reconstructed using an iterative mapping and assembly procedure. The mitogenome of *T. areolatus* (GenBank accession AP012026, Nakatani, Miya, Mabuchi, Saitoh, & Nishida, 2011) was initially used as a seed reference. For each individual, paired-end reads mapping to this reference were kept and used to generate a *de novo* assembly. All original quality-filtered reads were then remapped against the *de novo* mitogenome to refine the consensus sequence. This cycle of mapping, assembly, and remapping was repeated iteratively for each taxon until curated and internally consistent final mitogenomes were obtained. The entire procedure was conducted in Geneious Prime v2026.0.2.

Newly generated mitogenomes were annotated in MITOS2 (Donath et al., 2019) to delimit ribosomal RNA genes, protein-coding genes, and tRNA genes. Annotations were confirmed against available mitogenomes of Trichomycteridae and other Loricarioidei (see below) to ensure homology among the sequence blocks used in subsequent analyses.

### 2.2 Public data, molecular taxon sampling, and comparative morphological material

The final mitogenomic matrix included 14 terminals (Table 1): seven mitogenomes generated in this study, together with the following complete mitogenomes available in GenBank and mitogenomic consensus sequences assembled from publicly available high-throughput sequencing reads deposited in NCBI databases: *T. areolatus* (GenBank accession AP012026), *T. caliensis* (GenBank accession PP261638), *Hoplisoma paleatum* (GenBank accession NC_063781), *Astroblepus grixalvii* (GenBank accession PP266002), and *Hypostomus ancistroides* (GenBank accession NC_052710). In addition, RNA-seq reads from *T. areolatus* collected in River Choapa, corresponding to SRA accessions SRR3714095 and SRR3714093, were downloaded, and their mitogenomes were assembled and refined following the procedure described above. Technical details regarding data retrieval from GenBank, quality control, assembly, mapping, consensus-sequence generation, and mitogenome annotation are provided in Appendix 1.

**TABLE 1.**
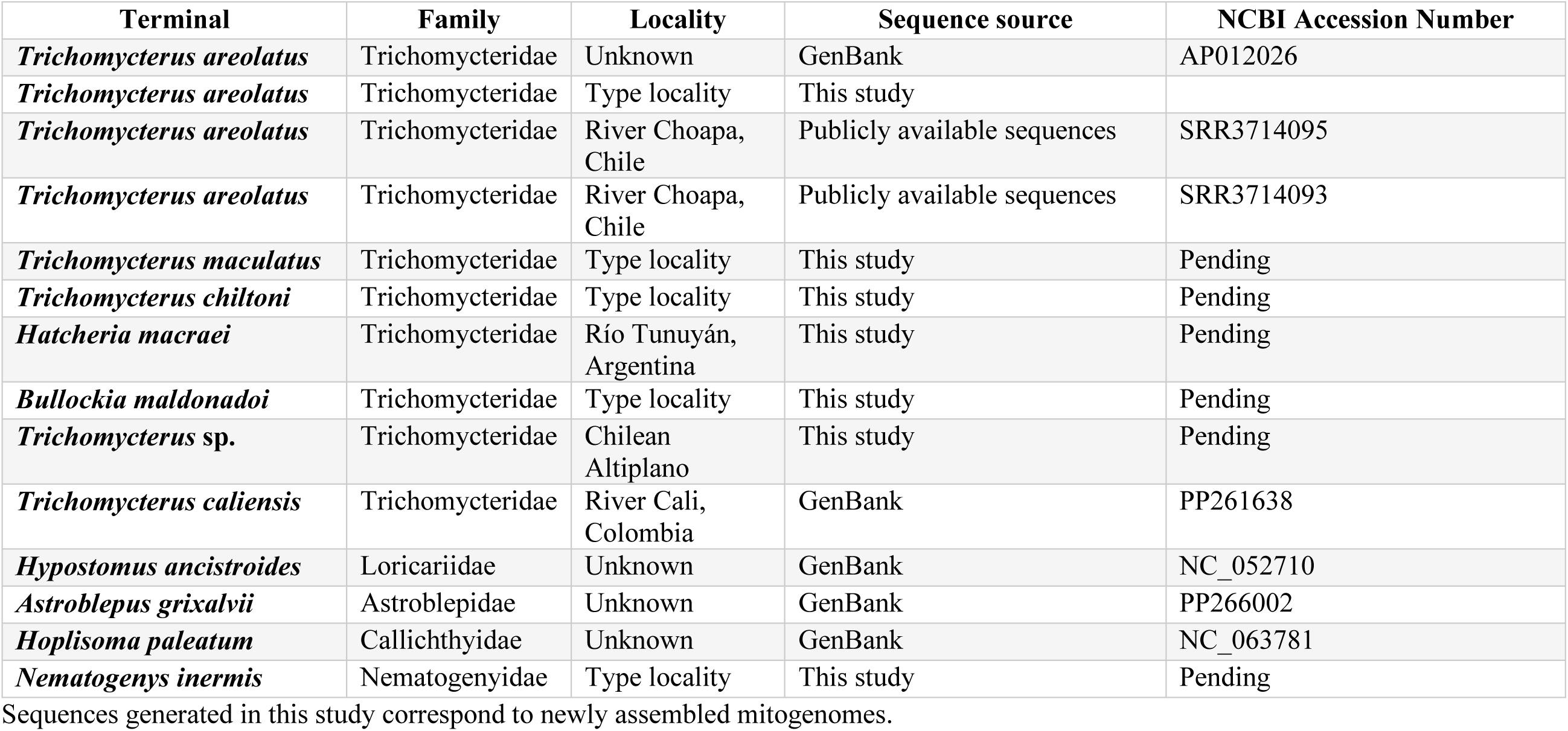
Taxon sampling, locality information and sequence origin for the mitogenomic dataset used in phylogenetic analyses.

Comparative morphological material included specimens assignable to *T. areolatus*, *T. maculatus*, *T. chiltoni*, *B. maldonadoi*, and *H. macraei*. This material was used to compare external characters, colouration patterns, and the configuration of the cephalic laterosensory system, with emphasis on the neurocranial canals in the supraorbital region (see below).

### 2.3 Mitogenomic alignment

The primary phylogenetic matrix included mitochondrial protein-coding genes, tRNA genes, and the ribosomal genes 12S rRNA and 16S rRNA. A preliminary alignment was generated in Geneious Prime v2026.1.2 to assess the suitability of the downloaded mitogenomic sequences and those assembled in this study. Each protein-coding gene was subsequently extracted, translated using the vertebrate mitochondrial genetic code, and examined to verify reading-frame conservation and the absence of internal stop codons. Genes encoded on the opposite strand were analysed in their coding orientation. Protein-coding genes were individually aligned using MAFFT v7.505 (Katoh & Standley, 2013), manually inspected, and concatenated while preserving codon positions. The tRNA genes were aligned and concatenated as a single independent block, whereas 12S rRNA and 16S rRNA were aligned and treated as separate partitions. The control region and other non-coding regions were excluded from the primary matrix.

The initial partitioning scheme (Chernomor, Von Haeseler, & Minh, 2016) treated each protein-coding gene separately by codon position, together with one partition for the concatenated tRNA genes and two additional partitions for 12S rRNA and 16S rRNA. Substitution-model selection and partition merging were performed in IQ-TREE v3.1.3 (Wong et al., 2026) using ModelFinder (Kalyaanamoorthy, Minh, Wong, Von Haeseler, & Jermiin, 2017), the MFP+MERGE option, and the Bayesian information criterion. The concatenated alignment and final partitioning scheme were used in all subsequent phylogenetic analyses.

### 2.4 Phylogenetic inference

Phylogenetic inference was performed under maximum likelihood in IQ-TREE v3.1.3 using the concatenated mitogenomic matrix and the partitioning scheme selected with ModelFinder under the merged-partition option. Node support was estimated using ultrafast bootstrap and SH-aLRT (Hoang, Chernomor, Von Haeseler, Minh, & Vinh, 2018), with nodes considered strongly supported when ultrafast bootstrap values were ≥95 and SH-aLRT values were ≥80. The tree was rooted using *Nematogenys inermis* (Genbank accession XXX), *Hoplisoma paleatum* (NC063781), *Hypostomus ancistroides* (NC052710), and *Astroblepus grixalvii* (PP266002) as outgroups. The resulting topology was used as the primary phylogenetic hypothesis for evaluating relationships among the Trichomycterinae lineages included in the study.

### 2.5 Tests of monophyly of *T. areolatus sensu lato*

The monophyly of *T. areolatus sensu lato* was evaluated in IQ-TREE v3.1.3 (Wong et al., 2026) using site-wise log-likelihoods and tree-comparison tests, including AU, KH, SH, weighted KH, weighted SH, and ELW tests with RELL resampling. The unconstrained maximum-likelihood topology was compared with an alternative topology inferred under a monophyly constraint comprising the terminals *T. areolatus* (AP012026), *T. areolatus* (type locality), *T. areolatus* (SRR3714095), *T. areolatus* (SRR3714093), and *T. maculatus* (type locality). Both topologies were estimated using the same mitogenomic matrix, partitioning scheme, substitution models, and outgroups. The hypothesis of monophyly of *T. areolatus sensu lato* was considered rejected when the AU test returned p < 0.05.

### 2.6 ML model-corrected patristic distances

Genetic distances among terminals were estimated as patristic distances from the maximum-likelihood tree obtained using the partitioned mitogenomic matrix. These distances were calculated as the sum of the branch lengths separating each pair of terminals and therefore represent divergences corrected according to the substitution models selected for the partitions used in the analysis. Comparisons were summarised for taxonomically relevant lineages, with emphasis on *T. areolatus sensu stricto*, *T. maculatus*, terminals historically assigned to *T. areolatus sensu lato*, *T. chiltoni*, *B. maldonadoi*, and *H. macraei*. Values were reported as substitutions per site and as percentage divergence corrected under the ML models.

### 2.7 Comparative morphological analysis: lateral-line system and colouration

Morphological comparisons focused on the cephalic lateral-line system and external colouration of the lineages recovered in the mitogenomic analyses, particularly those from the type locality of *T. areolatus*. The laterosensory system was examined in cleared-and-stained specimens and comparative osteological illustrations. Terminology for cephalic canals and pores followed Arratia and Huaquín (1995), consistent with previous studies of Trichomycteridae that adopted this nomenclature for the laterosensory system (Henschel et al., 2018). The analysis focused on the supraorbital canal, identifying the continuity or interruption of the canal and the presence or absence, as well as the positions of pores s1, s2, s3, and s6. For each specimen, we evaluated whether the supraorbital pores were connected by a continuous canal or represented by isolated canal segments. External colouration was assessed from photographs of live specimens or specimens photographed shortly after capture.

### 2.8 Integrative evaluation of taxonomic evidence

Taxonomic delimitation was evaluated using an integrative approach that considered the phylogenetic position of terminals, nodal support, monophyly tests, model-corrected genetic distances, characters of the cephalic laterosensory system, external colouration, and the geographic provenance of specimens. Taxonomic decisions were based on congruence among mitogenomic lineages, comparable morphological characters, and geographic distributions. Evidence was considered to support the recognition of taxonomically distinct lineages when they were recovered as independent phylogenetic units, exhibited genetic divergence comparable to that observed among recognised species, and displayed concordant morphological characters or geographic patterns. External colouration was treated as complementary evidence. Available names were interpreted according to their historical association with type localities and the available morphological and molecular evidence. Populations for which the evidence was insufficient to support formal species-level assignment were conservatively treated as *sensu lato*.

## 3. Results

### 3.1 Concatenated mitogenomic matrix and unconstrained phylogenetic hypothesis

The concatenated mitogenomic matrix included 14 terminals and 15 aligned blocks: the ribosomal genes 12S rRNA and 16S rRNA, 12 protein-coding genes, and a concatenated tRNA gene block. All blocks were represented in every terminal taxon, with no missing data in the final matrix. The concatenated alignment had a total length of 15,675 bp, of which 11,443 bp corresponded to protein-coding genes and 4,232 bp to ribosomal and tRNA genes.

Aligned blocks ranged in length from 168 bp for ATP8 to 1,829 bp for ND5. The ribosomal genes comprised 960 bp for 12S rRNA and 1,700 bp for 16S rRNA, whereas the concatenated tRNA block comprised 1,572 bp. This matrix was used to infer the unconstrained maximum-likelihood phylogenetic hypothesis (Figure 2).

**FIGURE 2.**
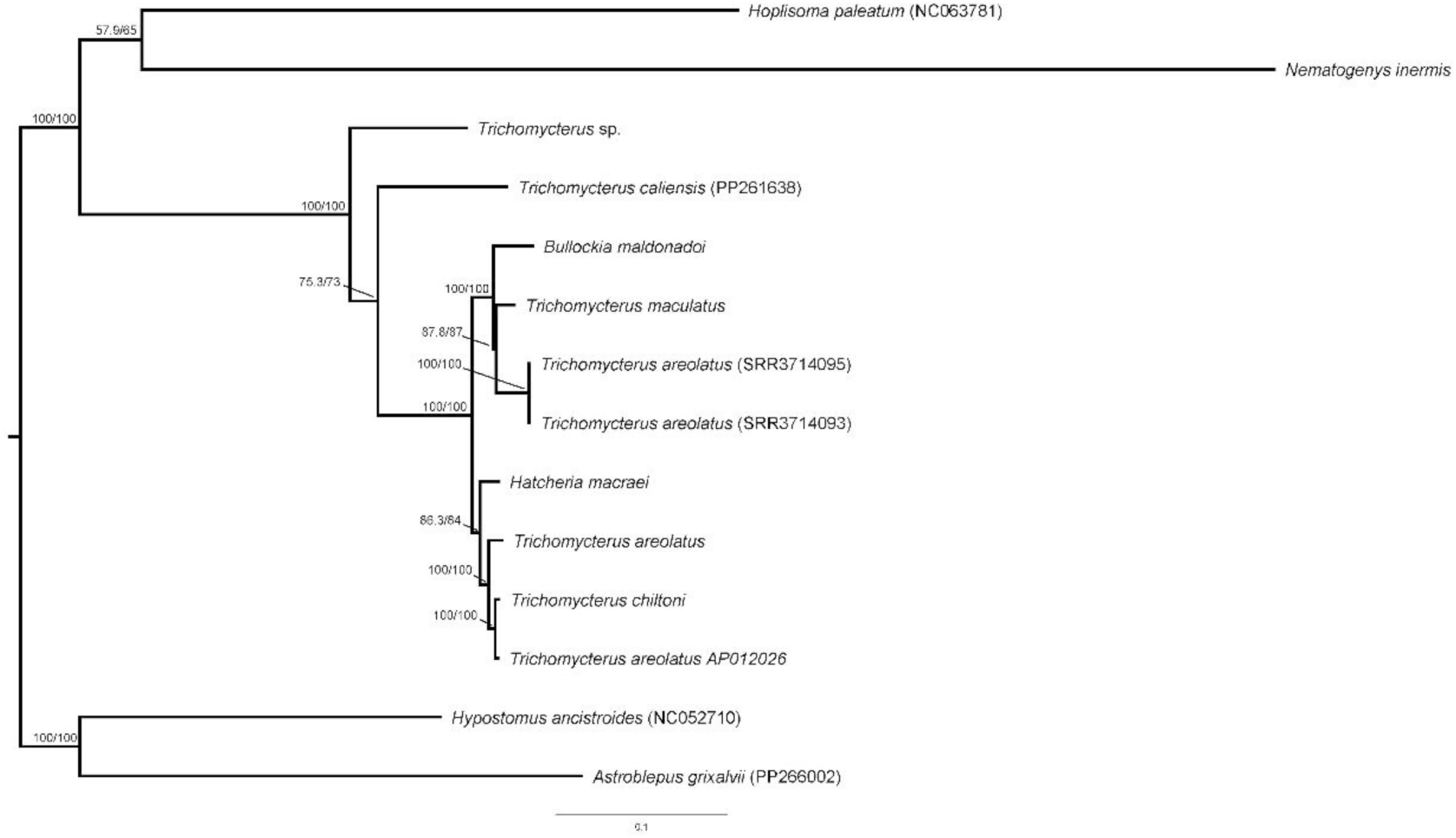
Maximum-likelihood phylogeny inferred from the concatenated mitogenomic matrix under an unconstrained topology. Node values indicate SH-aLRT support/ultrafast bootstrap support. Accession numbers are indicated in parentheses when available. The scale bar represents substitutions per site.

The ML tree recovered the Trichomycteridae included in the analysis as a well-supported clade. Within this clade, *Trichomycterus* sp. from the Chilean Altiplano was recovered as the earliest-diverging lineage relative to the remaining trichomycterines analysed, followed by *T. caliensis*. The remaining terminals formed a strongly supported clade comprising *B. maldonadoi*, *H. macraei*, *T. chiltoni*, *T. maculatus*, and the terminals assigned to *T. areolatus*.

The unconstrained hypothesis did not recover *T. areolatus sensu lato* as monophyletic (Figure 2). The two terminals obtained from RNA-seq reads assigned to *T. areolatus* (SRR3714093 and SRR3714095) formed a well-supported clade associated with *T. maculatus* and *B. maldonadoi*. In contrast, the *T. areolatus* terminal from the type locality generated in this study was recovered in a separate clade together with *H. macraei*, *T. chiltoni*, and the *T. areolatus* mitogenome AP012026. Within this latter clade, *T. chiltoni* was recovered as the sister lineage to *T. areolatus* AP012026, whereas the *T. areolatus* terminal from the type locality was recovered as sister to this pair.

### 3.2 Evaluation of the monophyly of *Trichomycterus areolatus sensu lato*

Comparison between the unconstrained maximum-likelihood tree and the alternative topology inferred under a monophyly constraint revealed substantial differences in log-likelihood, with the unconstrained topology showing a markedly higher likelihood than the constrained topology. In the analysis partitioned by codon position, the unconstrained tree had a log-likelihood of −68,618.93, whereas the constrained tree had a log-likelihood of −69,510.84, representing a difference of 891.91 log-likelihood units. The AU test rejected the constrained monophyly hypothesis (p = 4.16 × 10⁻⁶). The KH, SH, weighted KH, weighted SH, and ELW tests showed the same pattern, with values consistently supporting rejection of the constrained topology.

### 3.3 ML model-corrected patristic distances

Comparisons among terminals assigned to *T. areolatus sensu lato* showed heterogeneous values, consistent with their separation into more than one mitogenomic lineage (Table 2). The two consensus sequences derived from publicly available reads of *T. areolatus* from the River Choapa exhibited the lowest divergence from one another, whereas their comparisons with *T. areolatus* from the type locality and with *T. areolatus* AP012026 yielded higher values. *Trichomycterus maculatus* showed divergences of 3.1%–4.2% from terminals assigned to *T. areolatus sensu lato*, with values of 3.1% relative to the River Choapa consensus sequences and 4.1%–4.2% relative to *T. areolatus* AP012026 and *T. areolatus* from the type locality, respectively.

**TABLE 2.** Model-corrected patristic distances among focal trichomycterine terminals included in the *Trichomycterus areolatus* sensu lato comparison.

| **Taxon** | **1** | **2** | **3** | **4** | **5** | **6** | **7** | **8** |
| --- | --- | --- | --- | --- | --- | --- | --- | --- |
| **1. *Bullockia maldonadoi*** | — |  |  |  |  |  |  |  |
| **2. *Trichomycterus maculatus* (type locality)** | 3,6 | — |  |  |  |  |  |  |
| **3. *Trichomycterus areolatus* (SRR3714095)** | 4,5 | **3,1** | — |  |  |  |  |  |
| **4. *Trichomycterus areolatus* (SRR3714093)** | 4,5 | **3,1** | 0,1 | — |  |  |  |  |
| **5. *Hatcheria macraei*** | 5,1 | **4,0** | 4,9 | 4,9 | — |  |  |  |
| **6. *Trichomycterus areolatus* (type locality)** | **5,3** | **4,2** | **5,1** | **5,1** | **2,5** | — |  |  |
| **7. *Trichomycterus chiltoni*** | 5,2 | **4,1** | 5,0 | 5,0 | 2,4 | 1,5 | — |  |
| **8. *Trichomycterus areolatus* (AP012026)** | 5,2 | **4,1** | 4,9 | 5,0 | 2,3 | 1,5 | 0,6 | — |

Values below the diagonal represent percentages of divergence corrected under the ML model and estimated as patristic distances from the unconstrained maximum-likelihood tree. Taxa are ordered according to the focal clade recovered in the unconstrained ML topology.

Comparisons involving *T. chiltoni*, *B. maldonadoi*, and *H. macraei* provide a reference framework for assessing the magnitude of divergence among lineages currently treated within *Trichomycterus*. Overall, distances within the focal clade indicate that divergence between *T. maculatus* and the terminals assigned to *T. areolatus sensu lato* falls within the upper range observed among closely related nominal lineages of Trichomycterinae included in this analysis (Table 2). *Trichomycterus chiltoni* showed the lowest divergence from *T. areolatus* from the type locality and from terminal AP012026, and the highest divergence from *B. maldonadoi* (1.5%, 1.5%, and 5.2%, respectively). In contrast, *H. macraei* exhibited divergences of 4.0%–5.1% from *T. maculatus*, the River Choapa *T. areolatus* consensus sequences, and *B. maldonadoi*, whereas divergences from *T. areolatus* from the type locality and *T. chiltoni* ranged from 2.3% to 2.5% (Table 2).

### 3.4 Comparative morphological analysis: lateral-line system and colouration

Comparison of the cephalic laterosensory system revealed discrete differences between specimens assigned to *T. areolatus* and *T. maculatus* from the type locality. These differences were concentrated in the supraorbital region, particularly in the presence and arrangement of pores s1, s2, s3, and s6 (Figure 3).

**FIGURE 3.**
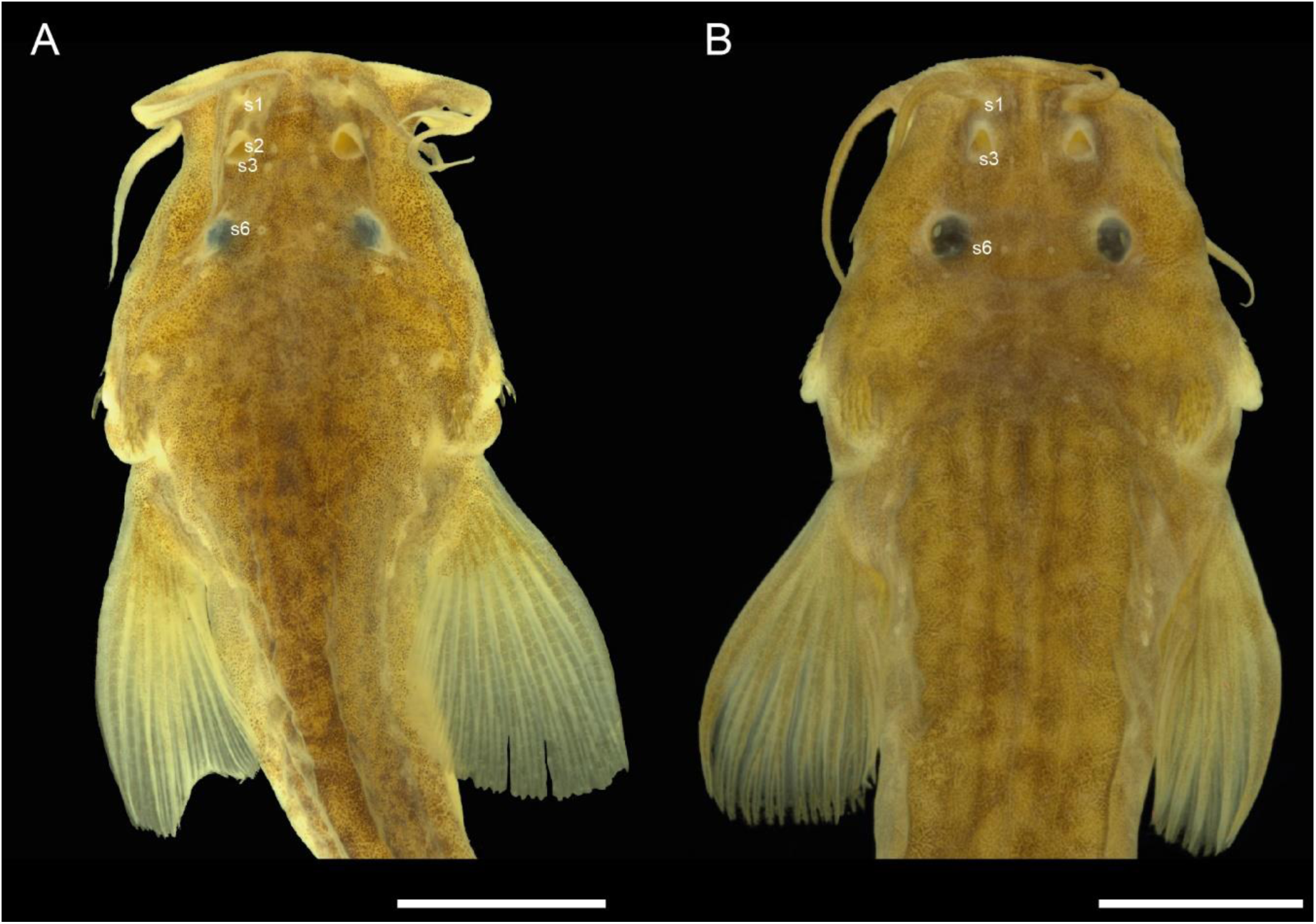
Dorsal view of the head and anterior body region of specimens showing the pores of the supraorbital cephalic laterosensory system. (A) *Trichomycterus areolatus* from the type locality, showing pores s1, s2, s3 and s6. (B) *Trichomycterus maculatus* from the type locality, showing pores s1, s3 and s6; pore s2 was not observed as a distinct pore. Abbreviations follow Arratia and Huaquín (1995). Scale bars: 1 cm.

In *T. areolatus*, the supraorbital canal exhibits pores s1, s2, and s3 in the anterior region of the neurocranium, together with pore s6 located posterior to the orbital margin. In dorsal view, pores s1– s3 are positioned close to one another and longitudinally aligned in the anterior ethmoidal–frontal region. The presence of s2 between s1 and s3 is evident both in the specimen fixed in ethanol (Figure 3) and in the comparative osteological illustration based on cleared-and-stained specimens (Figure 4).

**FIGURE 4.**
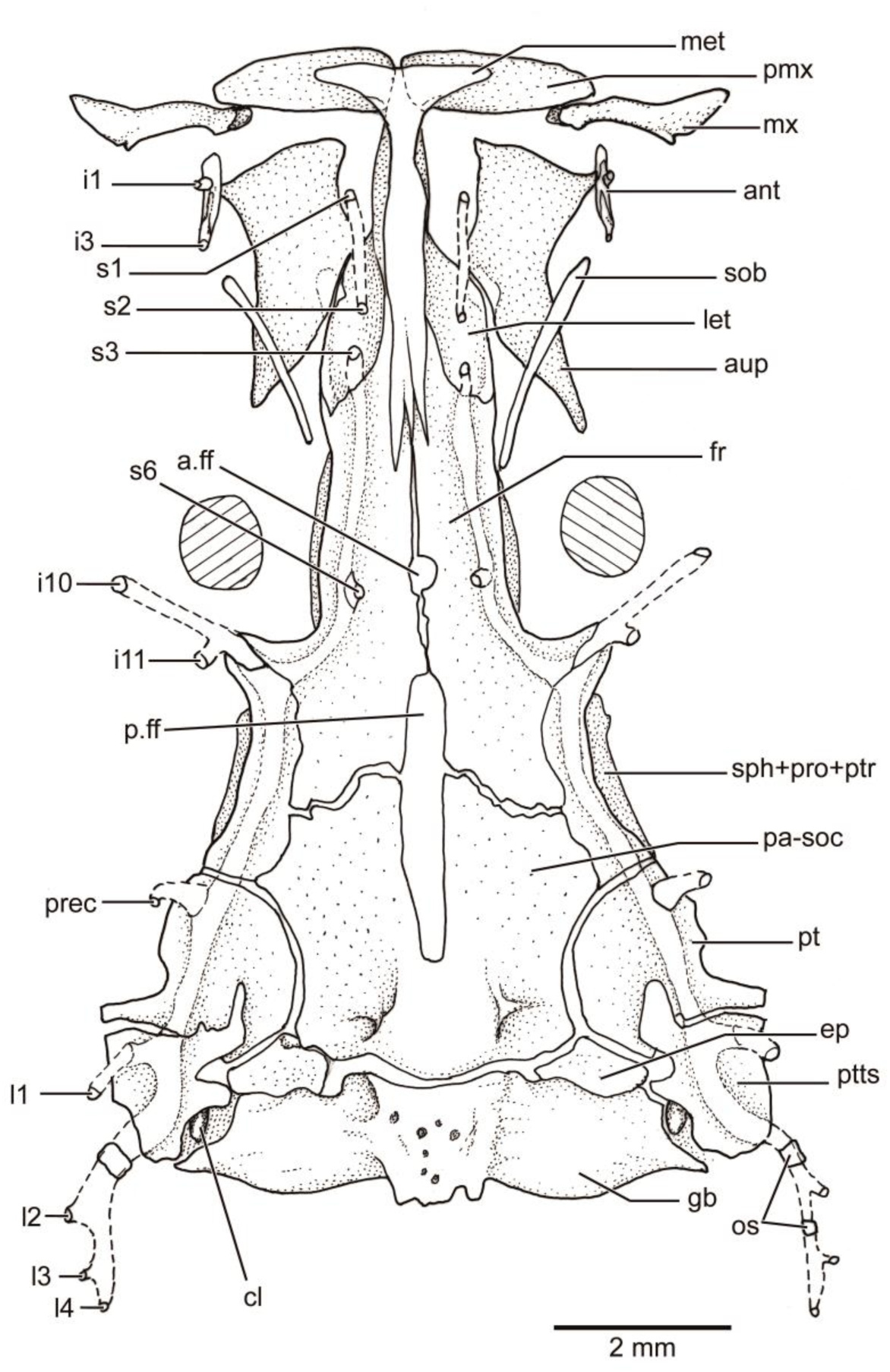
Dorsal view of the neurocranium and associated cephalic laterosensory canals of *Trichomycterus areolatus* from the type locality. The supraorbital series includes pores s1, s2, s3 and s6. Abbreviations: a.ff, anterior fontanelle; ant, antorbital; aup, autopalatine; cl, cleitrum; ep, epioccipital; fr, frontal; gb, gas bladder; i1–i3, infraorbital pores; i10–i11, infraorbital pores; let, lateral ethmoid; met, mesethmoid; mx, maxilla; os, ossicles; pa-soc, parieto-supraoccipital; p.ff, posterior fontanelle; pmx, premaxilla; prec, preopercular canal; pt, pterotic; ptts, posttemporo-supracleithrum; sph+pro+ptr, sphenotic-prootic-pterosphenoid complex ;s1–s6, supraorbital pores; sob, supraorbital. Scale bar = 2 mm.

In *T. maculatus*, the supraorbital pattern differs. Pores s1 and s3 are present, but no distinct s2 pore is observed between them. Pore s6 is present posteriorly, as in *T. areolatus* (Figure 3). In addition to differences in pore number, the continuity of the supraorbital canal also differs between the two taxa: in *T. maculatus*, the canal is continuous (Figure 5), whereas in *T. areolatus* it is interrupted (Figure 4). Consequently, the main differences between the two taxa involve the presence or absence of pore s2 and the organisation and continuity of the anterior segment of the supraorbital canal.

**FIGURE 5.**
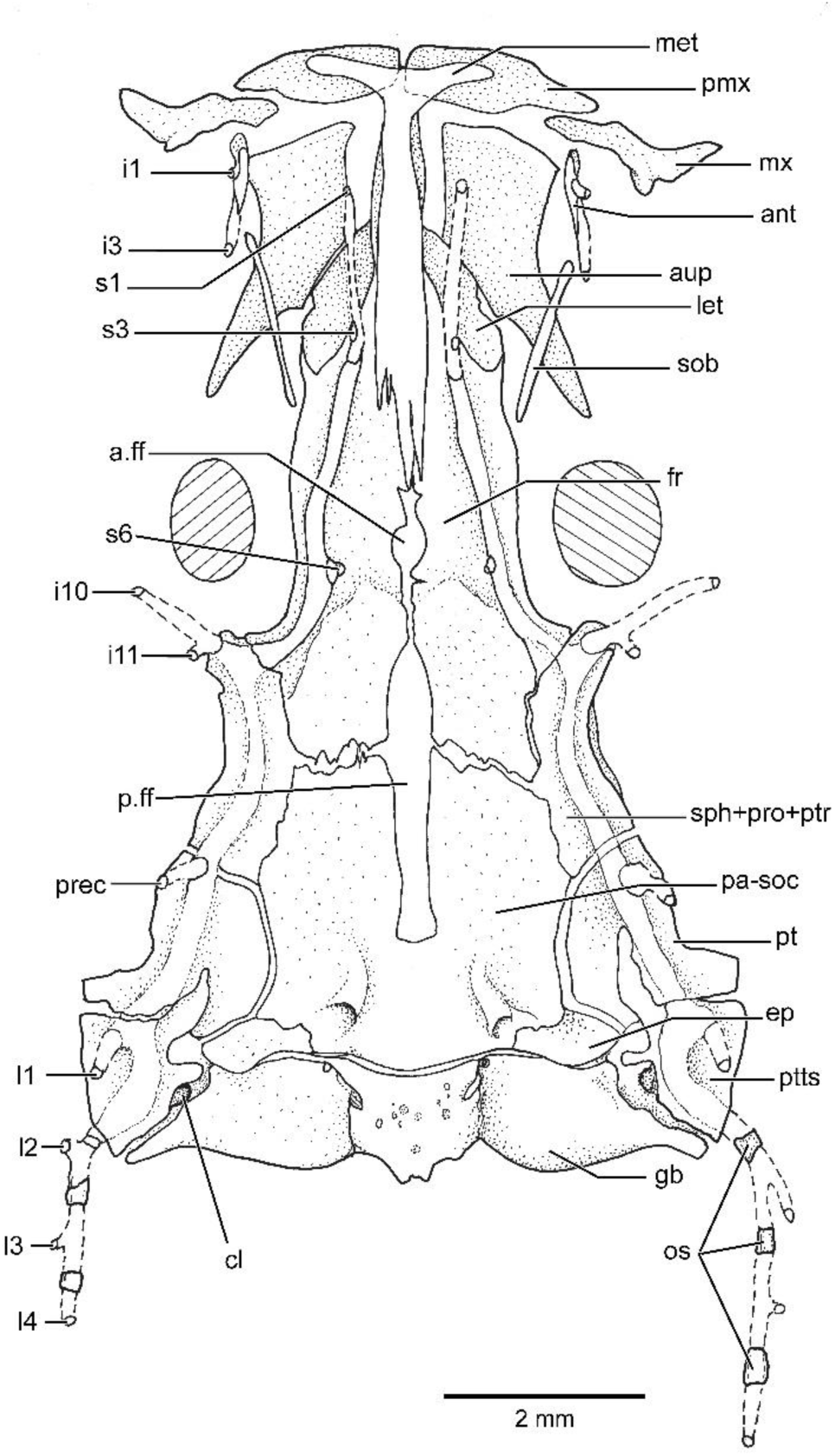
Dorsal view of the neurocranium and associated cephalic laterosensory canals of *Trichomycterus maculatus* from the type locality. The supraorbital series includes pores s1, s3 and s6; pore s2 was not observed as a distinct pore. Abbreviations: a.ff, anterior fontanelle; ant, antorbital; aup, autopalatine; cl, cleitrum; ep, epioccipital; fr, frontal; gb, gas bladder; i1–i3, infraorbital pores; i10–i11, infraorbital pores; let, lateral ethmoid; met, mesethmoid; mx, maxilla; os, ossicles; pa-soc, parieto-supraoccipital; p.ff, posterior fontanelle; pmx, premaxilla; prec, preopercular canal; pt, pterotic; ptts, posttemporo-supracleithrum; sph+pro+ptr, sphenotic-prootic-pterosphenoid complex;s1–s6, supraorbital pores; sob, supraorbital. Scale bar = 2 mm.

External colouration also differs between the specimens examined. *Trichomycterus areolatus* from the type locality exhibits more densely mottled dorsal pigmentation, with numerous small melanophores distributed over the head and anterior region of the body, together with the general presence of a longitudinal band extending along the body (Figure 6a–b). In *T. maculatus*, dorsal pigmentation is more homogeneous overall but conspicuously maculated, with large, dark, irregular blotches and greater contrast between the markings and the background colouration of the body (Figure 6c). These differences are observed in dorsal photographs of live specimens or specimens photographed shortly after fixation in ethanol and are presented as complementary evidence to the laterosensory characters.

**FIGURE 6.**
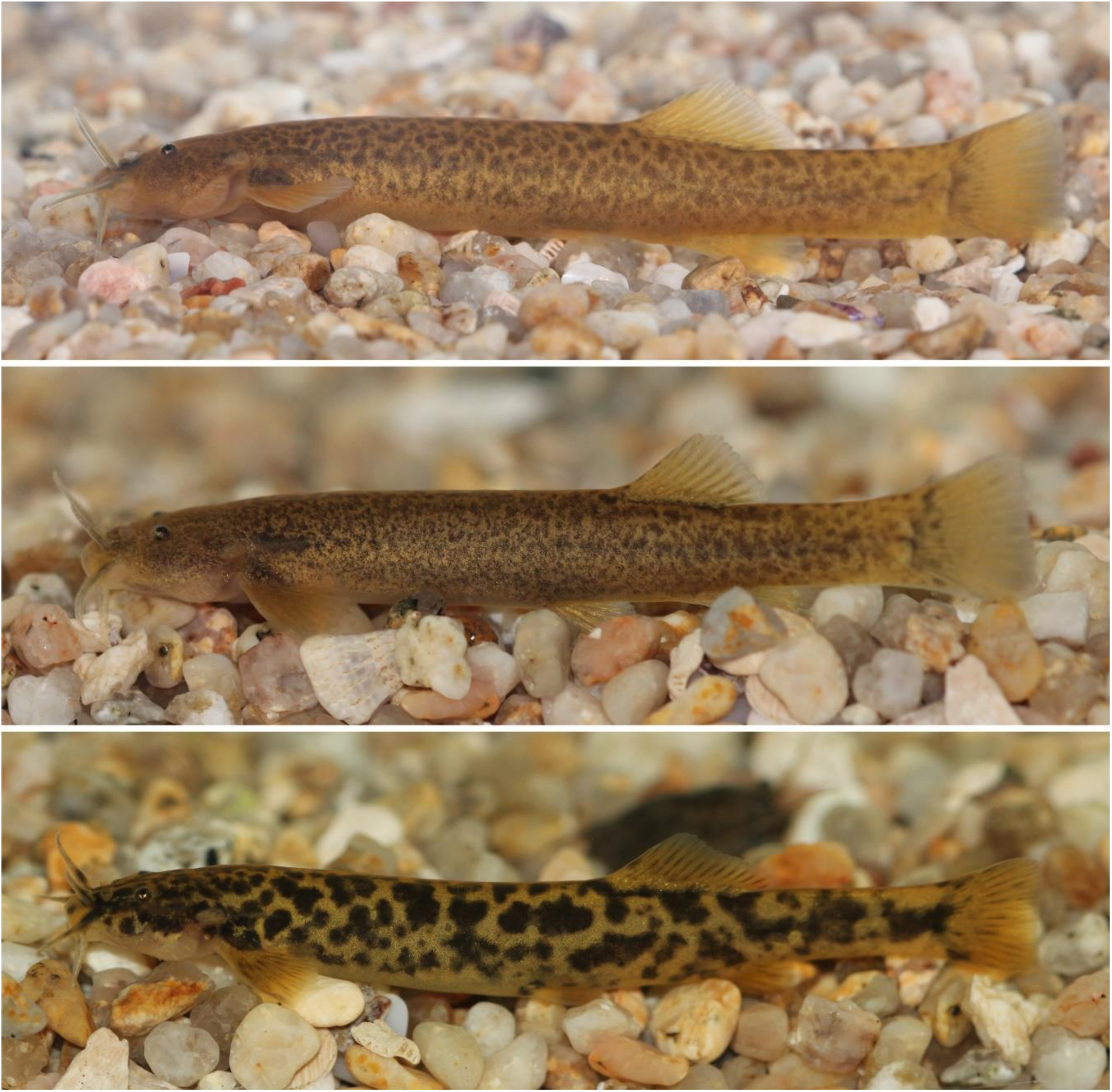
External colouration of live specimens of the focal *Trichomycterus* lineages. (a–b) *Trichomycterus areolatus* from type locality, showing a brown to olive-brown background colour with dense, fine mottling pattern distributed over the head, flank and caudal peduncle, without large, sharply delimited dark blotches. (c) *Trichomycterus maculatus* from type locality, showing a contrasting maculated pattern, with large, irregular dark blotches over a pale yellowish background, especially evident along the head, flank and caudal peduncle.

## 4. Discussion

The results of this study reveal four main patterns. First, the concatenated mitogenomic matrix recovered a topology in which the terminals assigned to *Trichomycterus areolatus sensu lato* did not form an exclusive clade. Second, the alternative hypothesis constraining *T. areolatus sensu lato* as monophyletic was rejected by all topology-comparison tests, including the AU test. Third, model-corrected patristic distances revealed substantial divergence among *T. maculatus*, *T. areolatus* from the type locality, and the other terminal taxa historically assigned to *T. areolatus*, with values comparable to those observed among nominal species within the focal clade. Finally, morphological comparisons revealed discrete differences in the cephalic laterosensory system, particularly in the continuity of the supraorbital canal and the number of pores in the supraorbital series, together with complementary differences in external colouration. These lines of evidence allow the taxonomic status of *T. maculatus* and the traditional circumscription of *T. areolatus* to be reassessed.

### 4.1 Non-monophyly of *Trichomycterus areolatus* sensu lato

The mitogenomic results reject the traditional interpretation of *T. areolatus* as a single, widely distributed species encompassing all focal lineages analysed in this study. These results were consistent across the codon-partitioned analysis (Figure 2), the gene-partitioned analysis, and the analysis based on complete mitogenomic sequences (results not shown). This finding agrees with previous molecular studies that identified strong geographic structure within *T. areolatus sensu lato*. The phylogeographic analyses of Unmack et al. (2009) and Barber et al. (2011) recovered differentiated mitochondrial and mito-nuclear lineages throughout the species’ distribution, whereas subsequent mitcochondrial and nuclear multilocus and phylogenomic studies placed *T. areolatus* in close association with other southern Andean trichomycterines, including *Bullockia maldonadoi*, *T. chiltoni*, and *Hatcheria macraei* (Fernandez et al., 2021; Ochoa et al., 2020, 2017). The principal difference in our study is that this evidence is not interpreted solely as phylogeographic structure, but is explicitly evaluated within a taxonomic framework by incorporating material associated with the type localities of *T. areolatus* and *T. maculatus*, populations provisionally assigned to *T. areolatus sensu lato*, and key taxa required to assess the non-monophyly of this species.

### 4.2 Mitogenomic divergence and comparison with valid species

The magnitude of model-corrected patristic distances supports the interpretation that *T. areolatus sensu lato* includes more than one taxonomic entity. The distance between *T. maculatus* and *T. areolatus* from the type locality was 4.2%, whereas distances between *T. maculatus* and other terminals assigned to *T. areolatus* ranged from 3.1% to 4.1%. These values overlap those observed among nominal taxa included in the focal clade, including comparisons involving *T. chiltoni*, *B. maldonadoi*, and *H. macraei* (Table 2). In Pangasiidae, genetic-distance analyses based on complete mitogenomic sequences have documented interspecific p-distances ranging from 4.24%–4.31% between *Pangasius mekongensis* and *P. pangasius*, and reaching 9.31% among other species within the family (Thuy Yen Duong et al., 2023). At the intraspecific level, that study reported divergences of approximately 0.07%–0.34% (Thuy Yen Duong et al., 2023).

In this context, it is noteworthy that *T. chiltoni*, a valid species within the focal clade, exhibited divergences of only 1.5% from *T. areolatus* from the type locality and the specimen AP012026, values that fall within the lower range observed among terminals of the clade and are substantially lower than those recorded between *T. maculatus* and the terminals assigned to *T. areolatus sensu lato* (3.1%-4.2%). The heterogeneity in genetic distances among terminals currently assigned to *T. areolatus sensu lato* –with values varying according to the pairwise comparison– suggests that this assemblage includes more than one differentiated mitogenomic lineage, whose taxonomic circumscription should be evaluated in future studies incorporating broader geographic and specimen sampling with valid species of the focal clade used as comparative reference points.

### 4.3 The cephalic laterosensory system as independent evidence

The cephalic laterosensory system provides an independent source of morphological evidence for distinguishing the focal lineages. In the material examined, *T. areolatus* from the type locality exhibits supraorbital pores s1, s2, s3, and s6, with the supraorbital canal interrupted between its nasal and frontal portions. In contrast, *T. maculatus* exhibits a continuous supraorbital canal, without a differentiated s2 pore between s1 and s3, while retaining pores s1, s3, and s6. This difference is discrete, topologically comparable, and observable in cleared-and-stained specimens, specimens preserved in EtOH, and live individuals.

The interpretation of these pores follows the nomenclature of Arratia and Huaquín (1995), who documented variation in the supraorbital canal among diplomystids and loricarioids, including Trichomycteridae. The presence or absence of pore s2 and the continuity of the supraorbital canal have subsequently been used as comparative and diagnostic characters in *Trichomycterus* and other trichomycterids (G. Arratia, 1998; DoNascimiento, Prada-Pedreros, & Guerrero-Kommritz, 2014, 2015; Henschel et al., 2018). Therefore, the congruence between mitogenomic differentiation and the configuration of the cephalic laterosensory system reinforces the interpretation of *T. maculatus* and *T. areolatus* from the type locality as evolutionarily differentiated lineages.

### 4.4 External colouration and distribution as complementary evidence

External colouration provides complementary support for the differentiation of the focal lineages. Specimens assigned to *T. areolatus* generally exhibit a finely mottled brown to olive-brown pattern, with dense but diffuse pigmentation over the head, flank, and caudal peduncle (Figure 6). In contrast, *T. maculatus* exhibits a more conspicuously maculated pattern, with large, irregular dark blotches against a lighter background (Figure 6). Because colouration may vary with ontogeny, environmental conditions, physiological state, and preservation, this character should be interpreted cautiously and not treated as diagnostic evidence in isolation. Its evidential value increases, however, when it is concordant with molecular and laterosensory evidence. Moreover, the maculated pattern observed in *T. maculatus* is consistent with the historical meaning of the specific epithet, further supporting its utility as a complementary external character.

The distributions of these species are illustrated in Figure 7. *Bullockia maldonadoi* occurs in rivers between approximately 36° and 38° S, whereas *T. maculatus* is restricted to Maipo and Rapel basins, between approximately 33° and 35° S. The SRR3714093 and SRR3714095 terminals associated with *T. maculatus* represent populations distributed from Huasco to Aconcagua basins, between approximately 28° and 33° S. *Hatcheria macraei* occurs in Chile only at approximately 45° S and, in Argentina, from this latitude northwards to approximately 32° S. *Trichomycterus areolatus sensu stricto* is restricted to Maipo and Mataquito basins, between approximately 33° and 35° S. The lineage represented by terminal AP012026 occurs from approximately 35° to 41° S and on Chiloé Island, whereas *T. chiltoni* is distributed between the Maule and Imperial basins, from approximately 35° to 38°50′ S.

### 4.5 Taxonomic implications

The congruence among phylogenetic position, monophyly tests, model-corrected genetic distances, laterosensory morphology, external colouration, and geographic provenance supports the recognition of *T. maculatus* as a species distinct from *T. areolatus*. Both names are historically associated with central Chile and with localities linked to the Santiago—Maipo area, although previous evidence had not explicitly assessed whether they represent two morphologically variable but distinguishable entities.

Accordingly to the results presented above, (1) *T. areolatus* should be interpreted in a restricted sense, associated with material from its type locality, characterised by the presence of supraorbital pores s1, s2, s3, and s6 and (2) *T. maculatus* should be revalidated for the lineage represented by material from its type locality, which exhibits the alternative supraorbital condition, lacking a differentiated s2 pore, together with a maculated external colour pattern, a morphology that seems to be unique among Chilean *Tricomycterus*. According to the present results, other Chilean populations historically assigned to *T. areolatus* should not be transferred automatically to either species without direct evidence. Until comparable molecular, morphological, and geographic data become available, populations south of Mataquito basin should be retained as *T. areolatus sensu lato*, whereas those north of Aconcagua basin should be retained as *T. maculatus sensu lato*.

This conservative treatment is particularly important because individuals from River Choapa and mitogenome AP012026 do not group with *T. areolatus* from the type locality, *T. maculatus* from the type locality, or *T. chiltoni*. Their phylogenetic positions suggest additional taxonomic complexity within this species group, but formal reassignment requires examination of voucher specimens, precise locality data, and diagnostic morphological characters. Therefore, this study resolves the status of *T. maculatus* and restricts the concept of *T. areolatus*, but highlights the need for a comprehensive taxonomic revision of the complex across its entire distribution, which is under study by the authors.

## 5. Conclusion

In summary, the broad concept of *Trichomycterus areolatus* has obscured a great taxonomic diversity within Chilean Trichomycterinae along an extense geographic distribution in central-south Chile. Complete mitogenomes show that *T. areolatus sensu lato* is not monophyletic, topology-comparison tests reject its constrained monophyly, and model-corrected patristic distances reveal divergences comparable to those observed among recognised species within the focal clade. These molecular results are concordant with discrete differences in the cephalic laterosensory system and complementary differences in external colouration. Taken together, the evidence supports the revalidation of *T. maculatus* and a restricted circumscription of *T. areolatus*.

